# A1 Adenosine Receptor Signaling Reduces *Streptococcus pneumoniae* Adherence to Pulmonary Epithelial Cells by Targeting Expression of Platelet-Activating Factor Receptor

**DOI:** 10.1101/670901

**Authors:** Manmeet Bhalla, Jun Hui Yeoh, Claire Lamneck, Sydney E. Herring, Essi Y. I. Tchalla, John M. Leong, Elsa N. Bou Ghanem

## Abstract

Extracellular adenosine production is crucial for host resistance against *Streptococcus pneumoniae* (pneumococcus) and is thought to affect antibacterial immune responses by neutrophils. However, whether extracellular adenosine alters direct host-pathogen interaction remains unexplored. An important determinant for lung infection by *S. pneumoniae* is its ability to adhere to the pulmonary epithelium. Here we explored whether extracellular adenosine can directly impact bacterial adherence to lung epithelial cells. We found signaling via A1 adenosine receptor significantly reduced the ability of pneumococci to bind human pulmonary epithelial cells. A1 receptor signaling blocked bacterial binding by reducing the expression of platelet-activating factor receptor, a host protein used by *S. pneumoniae* to adhere to host cells. *In vivo*, A1 was required for control of pneumococcal pneumonia as inhibiting it resulted in increased host susceptibility. As *S. pneumoniae* remain a leading cause of community-acquired pneumonia in the elderly, we explored the role of A1 in the age-driven susceptibility to infection. We found no difference in A1 pulmonary expression in young versus old mice. Strikingly, triggering A1 signaling boosted host resistance of old mice to *S. pneumoniae* pulmonary infection. This study demonstrates a novel mechanism by which extracellular adenosine modulates resistance to lung infection by targeting bacterial-host interactions.

## Introduction

*Streptococcus pneumoniae* (pneumococcus) typically reside asymptomatically in the nasopharynx of healthy individuals but can cause life-threatening invasive diseases resulting in more than 1 million deaths per year worldwide (Kadioglu, Weiser, Paton, & Andrew, 2008; van der Poll & Opal, 2009). The majority of invasive pneumococcal infections occur in elderly individuals ≥65 years of age (Chong & Street, 2008). As the number of elderly individuals is projected to double in the coming decades, reaching 2 billion by 2050, invasive pneumococcal infections pose a serious health concern and an economic burden which calls for novel interventions to fight this disease (Boe, Boule, & Kovacs, 2017).

Lung infection is the most common form of invasive pneumococcal disease (Centers for Disease Control and Prevention. 2016. Active Bacterial Core Surveillance Report, 2016) and the ability of *S. pneumoniae* to bind to and invade the airway epithelium is a crucial step for invasive infection (Kadioglu et al., 2008; van der Poll & Opal, 2009). *S. pneumoniae* express several factors that mediate binding to mammalian host cells and in fact many bacterial virulence elements with diverse functions also double as adherence factors (Kadioglu et al., 2008; van der Poll & Opal, 2009). This includes phosphorylcholine displayed on the bacterial cell wall that mimics moieties found on host platelet-activating factor (PAF) and allows the bacteria to bind to the host PAF receptor (Briles & Tomasz, 1973; Cundell, Gerard, Gerard, Idanpaan-Heikkila, & Tuomanen, 1995). Platelet-activating factor receptor (PAFR) is expressed on airway epithelial cells (Duitman et al., 2012; Ishizuka et al., 2001; Shivshankar, Boyd, Le Saux, Yeh, & Orihuela) and is upregulated during pneumococcal pneumonia (Duitman et al., 2012). PAFR mediates pneumococcal adhesion to airway epithelial cells (Cundell, Gerard, et al., 1995) and pharmacologically blocking this receptor decreased bacterial binding *in vitro* (Cundell, Gerard, et al., 1995; Shukla et al., 2016). Importantly blocking PAFR also attenuated *S. pneumoniae* infection in animal models and PAFR^-/-^ mice were more resistant to pneumococcal lung infection (Cundell, Gerard, et al., 1995; Rijneveld et al., 2004).

The extracellular adenosine pathway is an important regulator of host defense against *S. pneumoniae* infection (Bajgar & Dolezal, 2018; Bou Ghanem, Clark, Roggensack, et al., 2015). Upon cellular injury including that caused by infection, ATP is released from cells and converted to adenosine by the sequential action of two extracellular enzymes, CD39, which metabolizes ATP to AMP, and CD73 that de-phosphorylates AMP to adenosine (Hasko, Linden, Cronstein, & Pacher, 2008). Extracellular adenosine can then act as a signaling molecule by binding to and activating one of four G-protein coupled adenosine receptors, A1, A2A, A2B and A3 (Hasko et al., 2008). The enzymes and receptors of the extracellular adenosine pathway are expressed by lung epithelial cells (Burnstock, Brouns, Adriaensen, & Timmermans, 2012) and were shown to control acute lung injury (Hoegl et al., 2015), ciliary motility (Allen-Gipson et al., 2011), wound healing (Allen-Gipson, Wong, Spurzem, Sisson, & Wyatt, 2006), activation of ion channels (Factor et al., 2007), surfactant secretion (Gobran & Rooney, 1990), fibronectin release (Roman, Rivera, Roser-Page, Sitaraman, & Ritzenthaler, 2006), mucin expression (McNamara et al., 2004) and production of cytokines (Zhong, Wu, Belardinelli, & Zeng, 2006).

Extracellular adenosine is thought to regulate host resistance to lung infections by regulating recruitment and function of innate immune cells (Aeffner, Woods, & Davis, 2014, 2015; Barletta, Cagnina, Burdick, Linden, & Mehrad, 2012; Barletta, Ley, & Mehrad, 2012; Bou Ghanem, Clark, Roggensack, et al., 2015; Theatre et al., 2012). For example, we previously found that CD73 deficient mice had a thousand-fold more *S. pneumoniae* in their lungs as compared to wild type controls following lung challenge (Bou Ghanem, Clark, Roggensack, et al., 2015). However, those studies did not assess whether decreased adenosine functioned to enhance bacterial binding to lung epithelial cells. Here, we report that adenosine A1 receptor signaling inhibited binding of *S. pneumoniae* to human lung epithelial cells *in vitro* by reducing the expression of PAFR on infected host cells. Signaling via the A1 receptor was further required for control of bacterial numbers *in vivo* upon lung infection of mice. Importantly triggering A1 receptor signaling reversed the susceptibility of old mice to *S. pneumoniae* infection. This study identifies a novel mechanism by which adenosine regulates host resistance to bacterial pneumonia and demonstrates for the first time the feasibility of targeting this pathway to boost resistance of vulnerable hosts against pneumococcal pneumonia.

## Results

### Adenosine and AMP, but not ATP reduces binding of S. pneumoniae to H292 pulmonary epithelial cells

We previously found that extracellular adenosine was required for host resistance against *S. pneumoniae* lung infection (Bou Ghanem, Clark, Roggensack, et al., 2015). Mice lacking CD73, an enzyme required for extracellular adenosine production, had significantly higher bacterial burdens in their lungs as compared to wild type mice (Bou Ghanem, Clark, Roggensack, et al., 2015). Since a crucial step for establishing lung infection is the ability of *S. pneumoniae* to bind to the pulmonary epithelium (Kadioglu et al., 2008; van der Poll & Opal, 2009), we tested whether extracellular adenosine directly modulated pneumococcal binding to respiratory epithelial cells. To test that, we used H292 cells, a human lung epithelial cell line extensively used in modeling pulmonary bacterial host interaction (Bhowmick et al., 2013; Choi, Lee, Lee, Park, & Lee, 2008; R. T. Clark, Hope, Lopez-Fraga, Schiller, & Lo, 2009; Marks, Parameswaran, & Hakansson, 2012; Wagner et al., 2007; Yonker et al., 2017). H292 cells were treated with extracellular adenosine or vehicle control, infected with *S. pneumoniae* and the amount of bacteria that adhered to the host cells measured. We found that incubation with 100μM of extracellular adenosine, which is close to the alveolar concentration of adenosine in human subjects (Driver, Kukoly, Ali, & Mustafa, 1993), significantly decreased the amount of adherent bacteria that were attached to the lung epithelial cells by 3-fold (Fig 1A). This was not due to direct toxicity to the bacteria, as adenosine at the concentration we tested had no effect on bacterial growth as previously described (Bou Ghanem, Clark, Roggensack, et al., 2015) or bacterial viability within the timeframe of this assay (Fig S1).

**Fig 1.**
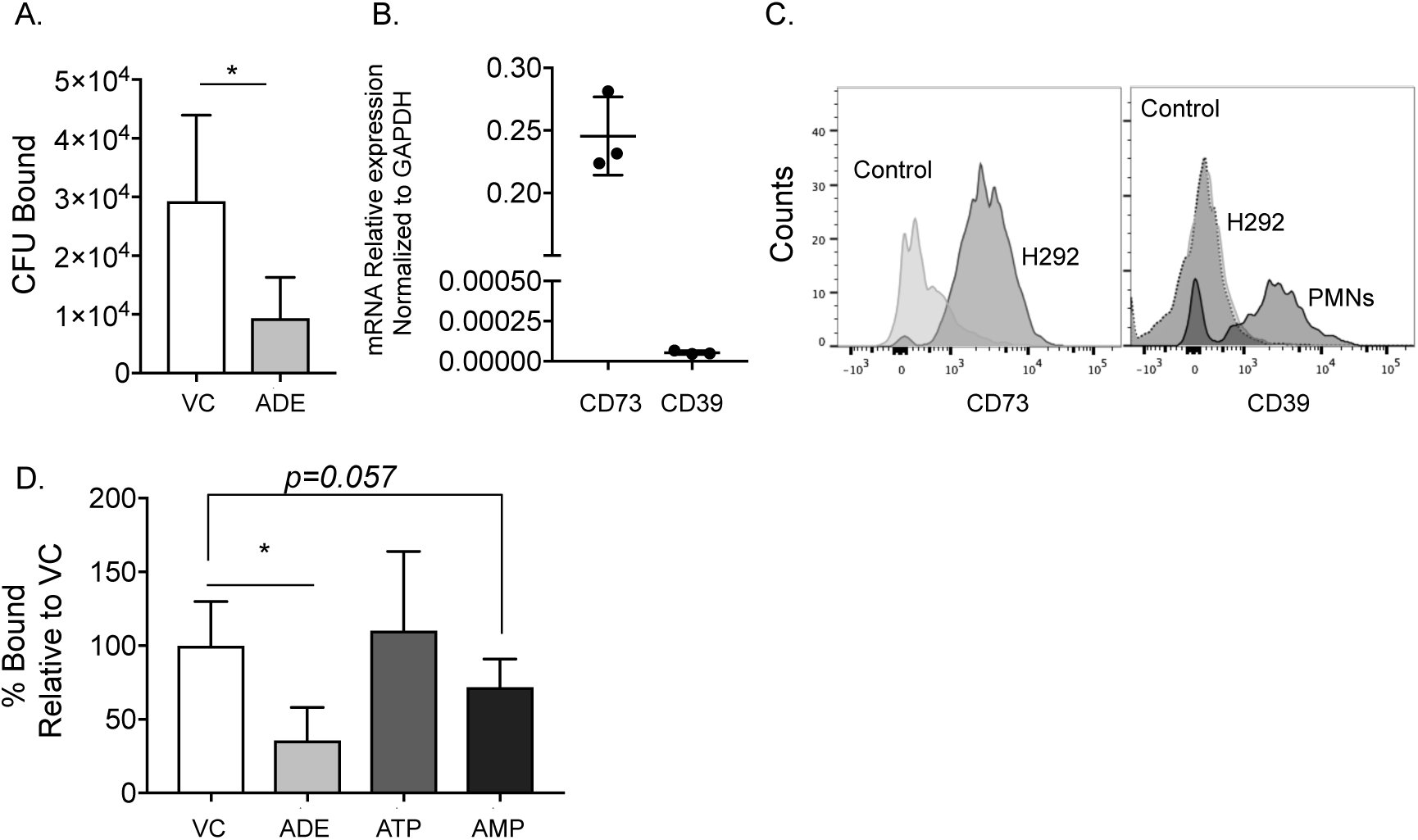
Extracellular adenosine reduces binding of *S. pneumoniae* to pulmonary epithelial cells. (A) H292 cells were treated with adenosine (ADE, 100µM) or vehicle control (VC) for 30 minutes and then infected with *S. pneumoniae* TIGR4 for 1 hour. The cells were then washed and the amount of bound bacteria were enumerated by plating on blood agar plates. (B) RNA was isolated from uninfected H292 cells and the expression of the indicated enzyme mRNA was measured by RT-qPCR and normalized to GAPDH. (C) Expression of CD73 (left panel) on H292 cells and CD39 on H292 cells and PMNs (right panel) was assessed by flow cytometry. Histograms shown are representative data from one of three separate experiments. (D) H292 cells were treated with either ADE, ATP, AMP (100µM) or vehicle control (VC) for 30 minutes and then infected with *S. pneumoniae* TIGR4 for 1 hour. The number of bound bacteria was enumerated by plating on blood agar plates. The percent of bacterial binding was then calculated relative to vehicle control treatment that was set to a 100. (A and D) Data shown are pooled from four separate experiments (n=4 biological replicates) where each condition was tested in triplicate (n=3 technical replicates) per experiment. Asterisks indicate significance calculated by (A) Student’s t-test and (B) one-way ANOVA followed by Tukey’s test.

Adenosine in the extracellular milieu is produced when ATP that has leaked from damaged cells is metabolized by two surface enzymes, CD39 that converts ATP to AMP, and CD73, an ecto-5’-nucleotidase that de-phosphorylates AMP to adenosine (Hasko et al., 2008). When we measured the expression of these enzymes in H292 cells, we found that mRNA levels of CD73 were readily detectable. However, CD39 mRNA levels were very low (Fig 1B). Consistent with that, CD73 protein was abundantly expressed on H292 cells, while CD39 was not detectable (Fig 1C). To confirm that the antibody we used for detection of CD39 was functional, we stained PMNs isolated from human donors as a positive control and were able to easily detect CD39 on those cells (Fig 1C).

Since H292 cells expressed CD73, but not CD39, we hypothesized that AMP will also be able to decrease binding of *S. pneumoniae* to these cells, since CD73 can convert it to adenosine, while ATP will have no effect since CD39 is not present to initiate its dephosphorylation into adenosine. When we performed the binding assays, we found that treatment of H292 cells with adenosine or AMP reduced the percentage of bacteria that bound when compared to vehicle control treatment (Fig 1D). However, we did not observe any significant effect on bacterial binding to host cells upon treatment with ATP. Taken together, these data suggest that extracellular adenosine production inhibits the ability of *S. pneumoniae* to bind to lung epithelial cells *in vitro*.

### Extracellular adenosine receptor signaling is required for reduction of S. pneumoniae binding to pulmonary epithelial cells

Extracellular adenosine can directly alter bacterial virulence (Crane & Shulgina, 2009; Kao et al., 2017) and also signal in the mammalian host via four known receptors (Hasko et al., 2008). Therefore we tested whether adenosine was altering bacterial binding by acting on the bacteria, the host or both. Capsule expression regulates the ability of *S. pneumoniae* to bind host cells (Cundell, Weiser, Shen, Young, & Tuomanen, 1995; Hammerschmidt et al., 2005; Sanchez, Hinojosa, et al., 2011; Sanchez, Kumar, et al., 2011; Trzcinski et al., 2015). However, treatment of *S. pneumoniae* with adenosine had no effect on expression of the capsular polysaccharide (Fig S2A). As expected capsular deficient (Δ*cps)* bacteria bound significantly better than wild type (WT) bacteria but adenosine treatment of H292 cells still reduced binding of Δ*cps S. pneumoniae* by 3-fold, comparable to its effect on WT bacteria (Fig S2B). Further, when we pre-treated *S. pneumoniae* only with adenosine, we saw no consistent decrease in bacterial binding (Fig 2). This suggests that adenosine was not mediating its effect through acting directly on the bacteria.

**Fig 2.**
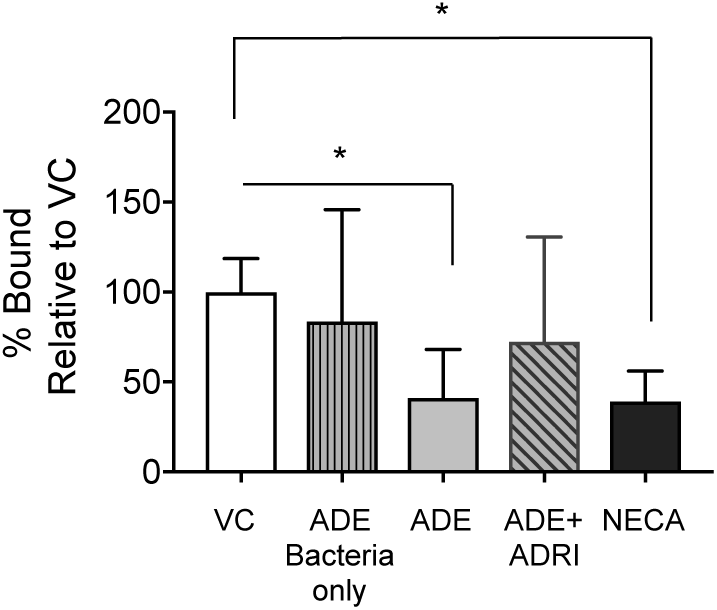
Signaling via extracellular adenosine receptors is required for reduction of *S. pneumoniae* binding to pulmonary epithelial cells. H292 cells were treated with either 100µM ADE, 100µM ADE in the presence of 1µM the pan adenosine receptors inhibitor (ADRI) CGS 15943, 1µM the pan adenosine receptors agonist 5’-*N*-Ethylcarboxamidoadenosine (NECA) or vehicle control (VC) for 30 minutes and then infected with *S. pneumoniae* TIGR4 for 1 hour. The number of bound bacteria was enumerated by plating on blood agar plates and the percent bacterial binding calculated relative to vehicle control. Data shown are pooled from three separate experiments where each condition was tested in triplicate. Asterisks indicate significance calculated by one-way ANOVA followed by Tukey’s test.

To test if adenosine was mediating its effect by acting on host cells, we blocked adenosine receptor signaling using a pan-adenosine receptor inhibitor CGS 15943. We found that the ability of adenosine to blunt bacterial binding to H292 cells was abrogated when adenosine receptors signaling was blocked (Fig 2). Further, treatment of H292 cells with NECA, a pan adenosine receptors agonist, mimicked the effects of adenosine, where bacterial binding was reduced to 39% compared to the vehicle treated controls (Fig 2). The drugs had no direct effect on bacterial viability as previously shown (Bou Ghanem, Clark, Roggensack, et al., 2015). These findings suggest that extracellular adenosine signaling in mammalian hosts inhibits the ability of *S. pneumoniae* to bind to lung epithelial cells.

### H292 cells express extracellular adenosine receptors

There are currently four known adenosine receptors A1, A2A, A2B and A3. First, we wanted to determine which adenosine receptors are expressed by H292 cells and whether their expression is affected by bacterial infection. To do so, we measured expression of adenosine receptors mRNA as is commonly done in most studies (Factor et al., 2007; Giacomelli et al., 2018; Roman et al., 2006; Zhong et al., 2006). mRNA for A1, A2A and A2B receptors was readily detectable, while A3 mRNA was very low (Fig 3A). Upon infection with *S. pneumoniae*, we saw a slight but not significant increase in mRNA levels for A1, A2A and A2B with no change in mRNA levels of A3 as compared to uninfected controls (Fig 3B). We then determined protein levels by flow cytometry and found that H292 cells expressed all four adenosine receptors and that A2A was more highly expressed on these cells compared to the other three receptors (Fig 3C). Adenosine receptor protein levels were not altered upon infection of H292 cells with *S. pneumoniae* (Fig 3C). We further confirmed our results with western blots and found that similar to flow cytometry, all 4 adenosine receptors were detected and *S. pneumoniae* infection slightly (but not significantly) increased levels of A1 by 1.4-fold but did not alter the expression of the other receptors (Fig 3D-E). These results demonstrate that receptors for extracellular adenosine are expressed in H292 pulmonary epithelial cells and that infection does not significantly alter their levels.

**Fig 3.**
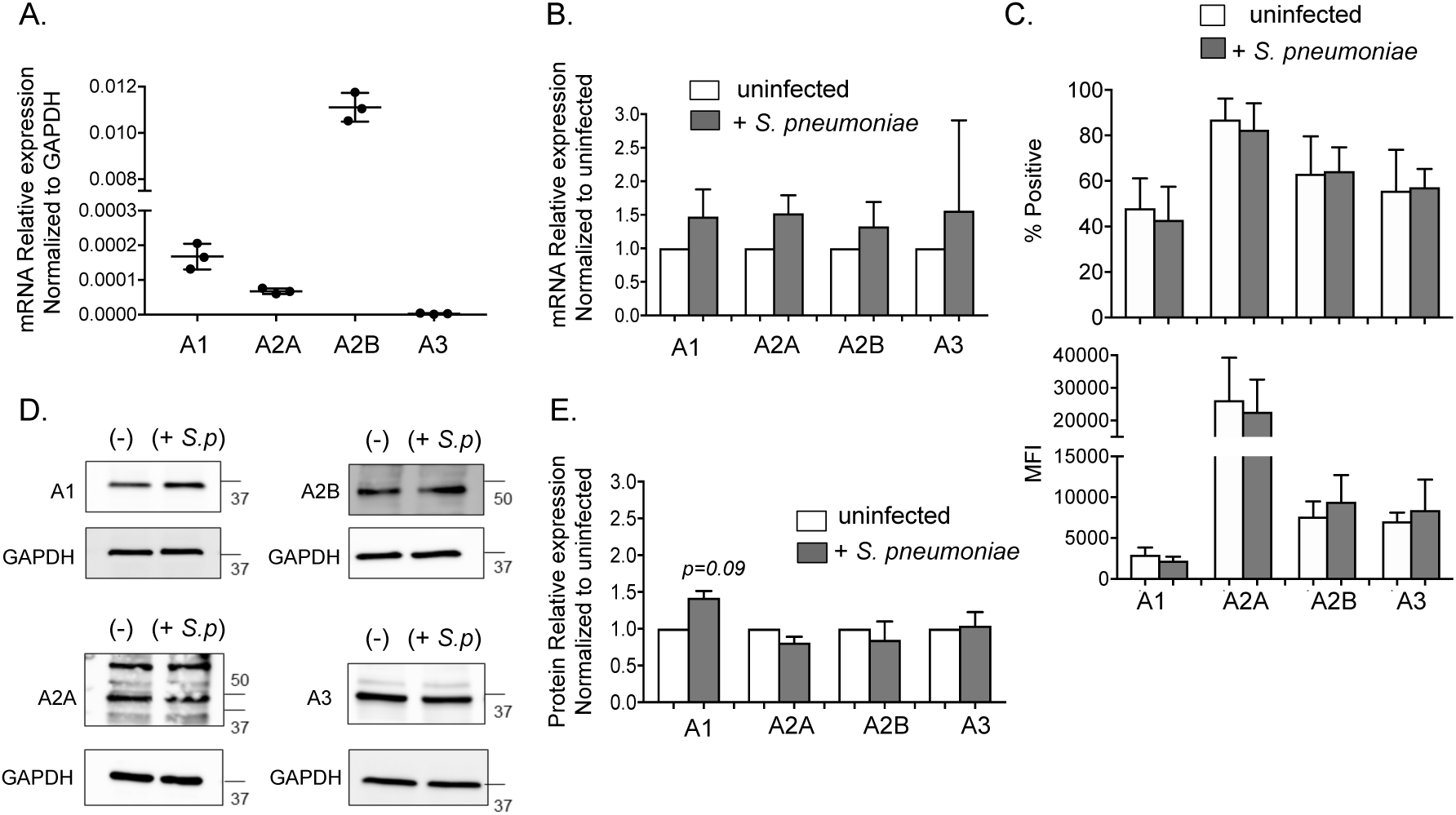
Expression of adenosine receptors on H292 cells. (A) RNA was isolated from uninfected H292 cells and the expression of the indicated adenosine receptor mRNA was measured by RT-qPCR and normalized to GAPDH. (B) RNA was isolated from H292 cells mock treated (uninfected) or challenged with *S. pneumoniae* TIGR4 MOI of 2 for 1 hour and the relative change in expression of the indicated adenosine receptor mRNA upon infection was calculated by the ΔΔCT method. Data shown are pooled from three separate experiments. H292 cells were infected with *S. pneumoniae* TIGR4 MOI of 2 for 1 hour or mock treated with PBS (uninfected). (C) Expression of adenosine receptors was assessed by flow cytometry. The percentage of cells expressing the receptors was then measured. Data shown are pooled from three separate experiments. (D) Expression of adenosine receptors was assessed by Western Blotting. One representative blot for each receptor is shown. (E) Western Blot band intensities were analyzed using ImageJ and expression of infected sample relative to uninfected controls is shown. Data presented are pooled from two experiments. (B and E) Adenosine receptor mRNA or protein levels upon infection were not significantly different from 1 by one-sample t-test.

### A1 receptor signaling reduces binding of S. pneumoniae to pulmonary epithelial cells

Next, we wanted to identify which adenosine receptor(s) was required for inhibiting bacterial binding to lung epithelial cells. These receptors bind to adenosine with different affinities where the higher affinity receptors A1 and A3 have an EC_50_ <0.5μM whereas the intermediate affinity A2A has an EC_50_ >0.6μM and the low affinity A2B an EC_50_ between 16-64μM (Hasko et al., 2008). To probe which receptor(s) is involved, we did a dose response, testing the effect of 100, 10, 1 and 0.1μM of adenosine on bacterial binding to H292 cells. We found that even lower concentrations of adenosine at 10 and 1μM were as equally efficient at decreasing adherence of *S. pneumoniae* as 100μM, where bacterial binding was reduced to 35% relative to vehicle treated controls (Fig 4). These results suggested that the higher affinity adenosine receptors A1, A2A or A3 but not the low affinity receptor A2B were mediating the effect on bacterial binding observed.

**Fig 4.**
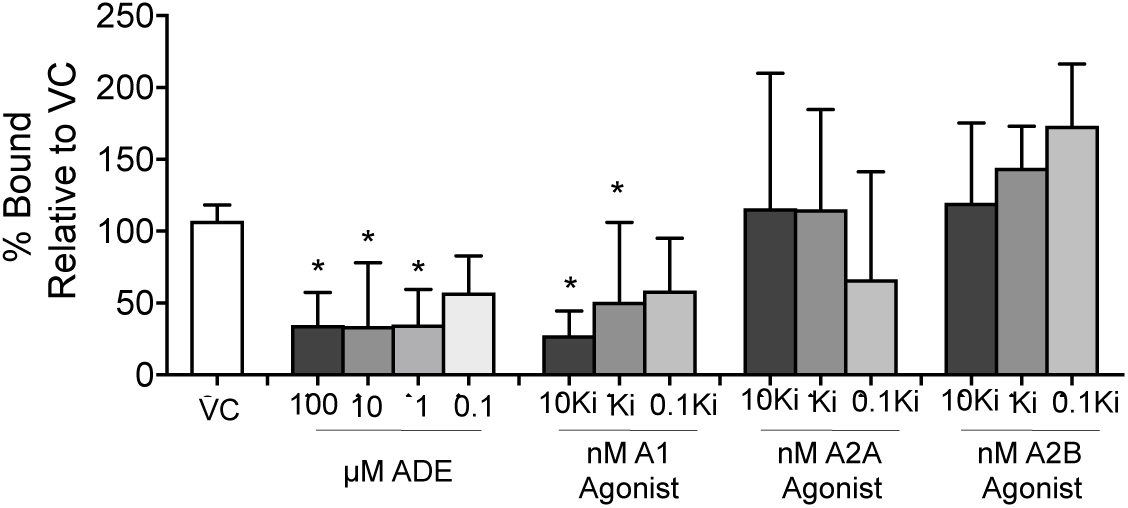
Signaling via A1 receptor is required for reduces binding of *S. pneumoniae in vitro* and *in vivo.* (A) H292 cells were treated for 30 minutes with either the indicated concentrations of ADE, vehicle control (VC) or at the following concentrations of A1 agonist 2-Chloro-N6-cyclopentyladenosine (10, 1, 0.1 nM), A2A agonist CGS21680 (270, 27, 2.7nM) or A2B agonist BAY 60-6583 (200, 20, 2nM) corresponding to the approximate 10xKi, Ki or 0.1Ki for each receptor as indicated on the graph. The cells were then infected with *S. pneumoniae* TIGR4 for 1 hour. The number of bound bacteria was determined by plating on blood agar plates and the percent bacterial binding calculated relative to vehicle control treatment. Data shown are pooled from three separate experiments where each condition was tested in triplicate. Asterisks indicate significant differences as compared to VC calculated by one-way ANOVA followed by Tukey’s test.

To pinpoint the exact receptor involved, we treated H292 cells with specific agonists for each of the different receptors. Since mRNA levels of A3 (Fig 3A) were barely detectable, we decided to proceed with testing the role of the other three receptors. Consistent with the results of adenosine doses, agonism of A2B receptor on H292 cells did not result in inhibition of *S. pneumoniae* binding to these pulmonary epithelial cells and the percentage of bacteria bound to treated cells were not statistically different than those bound to control treated cells (Fig 4). Similarly, treatment with the A2A receptor agonist did not alter bacterial binding to H292 cells (Fig 4). In contrast treatment with the A1 receptor agonist significantly reduced bacterial binding to H292 cells in a dose-dependent manner (Fig 4). We observed a 3-fold and 2-fold reduction in the amount of bound bacteria in comparison to control treated cells when cells were treated with 10 or 1 nM (corresponding to respectively 10x and 1x the drug’s reported Ki (Fig 4). None of the adenosine receptor agonists had a direct effect on bacterial viability (Fig S1). These findings suggest that specific signaling via the A1 receptor reduces binding of *S. pneumoniae* to pulmonary epithelial cells.

### A1 receptor signaling reduces expression of PAFR on infected pulmonary epithelial cells

Next we wanted to identify the mechanisms by which adenosine regulated bacterial binding to lung epithelial cells. *S. pneumoniae* is known to co-opt several well-characterized host proteins to adhere to mammalian cells including the Polymeric Immunoglobulin Receptor (PIGR) (Zhang et al., 2000) and PAFR (Cundell, Gerard, et al., 1995). Therefore, we examined the effect of adenosine on the expression of these host proteins by flow cytometry at baseline and during infection. As a control, we also examined the expression of ICAM-1 a host protein whose expression is upregulated during infection but is not thought to be a receptor for *S. pneumoniae* (Murdoch, Read, Zhang, & Finn, 2002). We found that consistent with previous studies, infection of H292 lung epithelial cells by *S. pneumoniae* increased the expression of all three proteins we examined (Fig 5A-C). Treatment of H292 cells with adenosine had no significant effect on the expression of ICAM-1 or PIGR (Fig 5A-B). In contrast treatment of the epithelial cells with adenosine or A1 receptor agonist prevented upregulation of PAFR in response to infection and *S. pneumoniae* infected H292 cells treated with these compounds had significantly lower expression of PAFR as compared to vehicle treated controls (Fig 5C).

**Fig 5.**
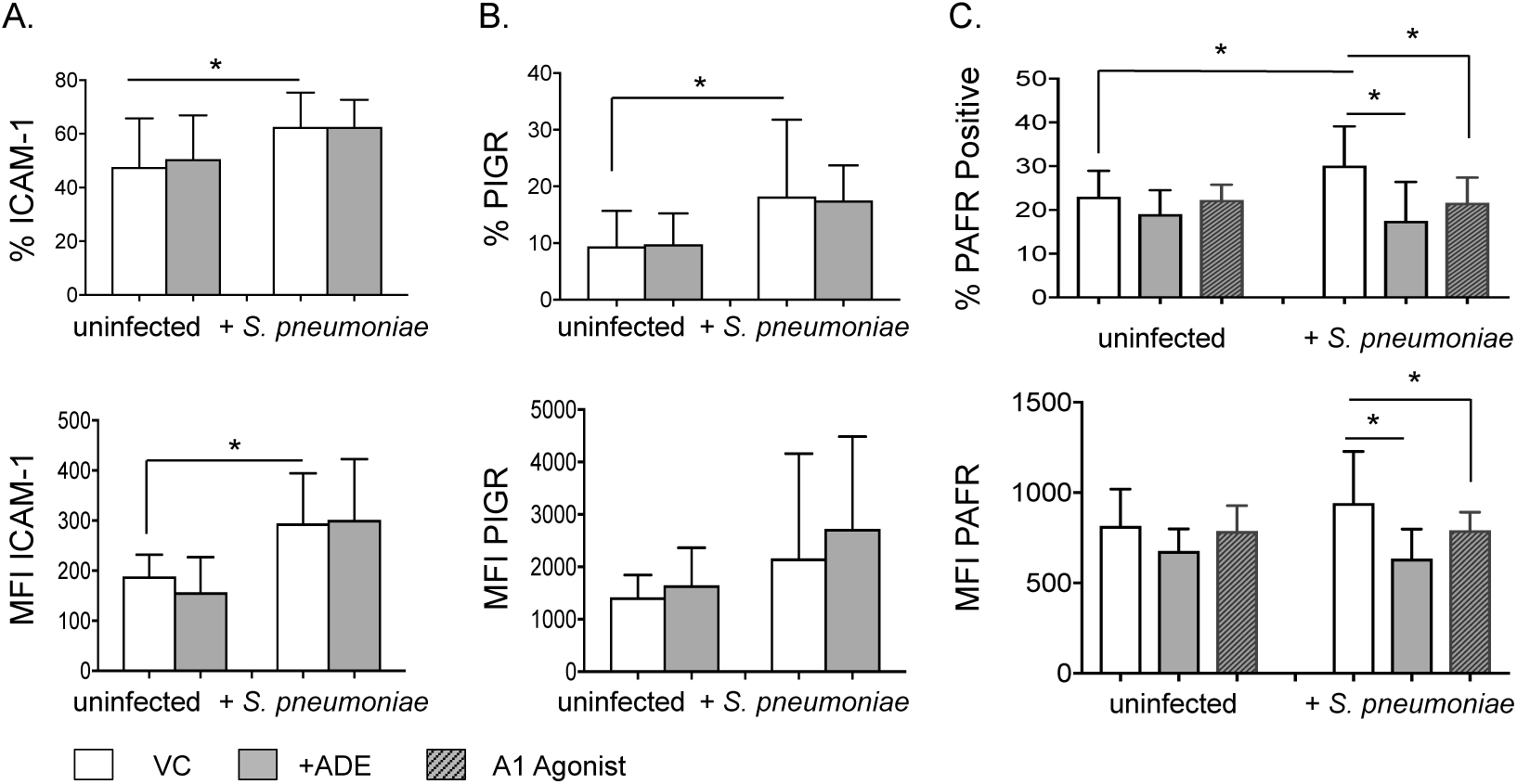
Signaling via A1 receptor reduces expression of PAFR on H292 cells. H292 cells were treated for 30 minutes with vehicle control (VC), ADE (100µM), or A1 agonist 2-Chloro-N6-cyclopentyladenosine (10nM) as indicated. The cells were then infected with *S. pneumoniae* TIGR4 MOI of 2 or mock treated (uninfected) for 1 hour. (A-C) Expression of ICAM-1, PIGR and PAFR was assessed by flow cytometry. The Mean florescent intensities (MFI) as well as the percentage of cells expressing the indicated ligands were then measured. (A-C) Data shown are pooled from three separate experiments where each condition was tested in triplicate. Asterisks indicate significant differences calculated by one-way ANOVA followed by Tukey’s test.

To test if adenosine was mediating its effect through decreasing expression of PAFR, we blocked PAFR signaling using the specific inhibitor Apafant. We found that as expected (Cundell, Gerard, et al., 1995; Shukla et al., 2016), inhibition of PAFR signaling significantly reduced the ability of *S. pneumoniae* to bind to H292 cells (Fig 6). Addition of the A1 receptor agonists or adenosine did not further decreased bacterial binding in the presence of the PAFR inhibitor (Fig 6). These findings suggest that A1 adenosine receptor signaling reduces bacterial binding to lung epithelial cells by targeting PAFR receptor.

**Fig 6.**
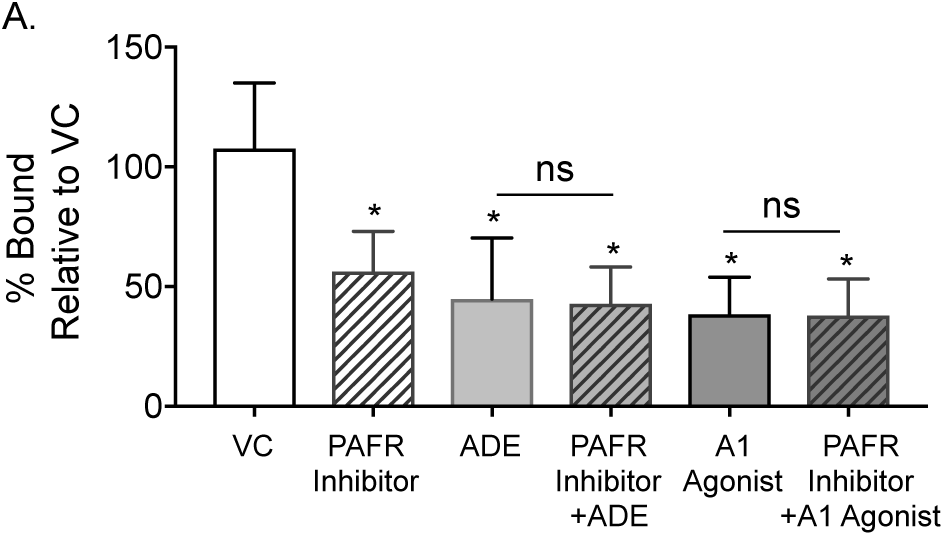
A1 receptor signaling does not further decrease bacterial binding when PAFR is inhibited. H292 cells were treated for 30 minutes with the indicated vehicle control (VC), ADE (100µM), A1 agonist 2-Chloro-N6-cyclopentyladenosine (10nM) or PAFR inhibitor (40nM). The cells were then infected with *S. pneumoniae* TIGR4 MOI of 20 for 1 hour. The number of bound bacteria was enumerated by plating on blood agar plates and the percent bacterial binding calculated relative to vehicle control. Data shown are pooled from four separate experiments where each condition was tested in triplicate. Asterisks indicate significant differences vs. vehicle control treated cells calculated by one-way ANOVA followed by Tukey’s test.

### A1 receptor signaling is required for host resistance to S. pneumoniae lung infection

We next wanted to confirm the relevance of our findings during lung infection *in vivo* using a mouse model. First we checked for the expression of adenosine receptors in the lungs of mice and found that consistent with previous reports (Factor et al., 2007), A1 and A2A were highly expressed, while less A2B and A3 was detected (Fig 7A-B). As we observed *in vitro*, *S*. *pneumoniae* infection did not affect the expression of adenosine receptors in mouse lungs (Fig 7A-B).

**Fig 7.**
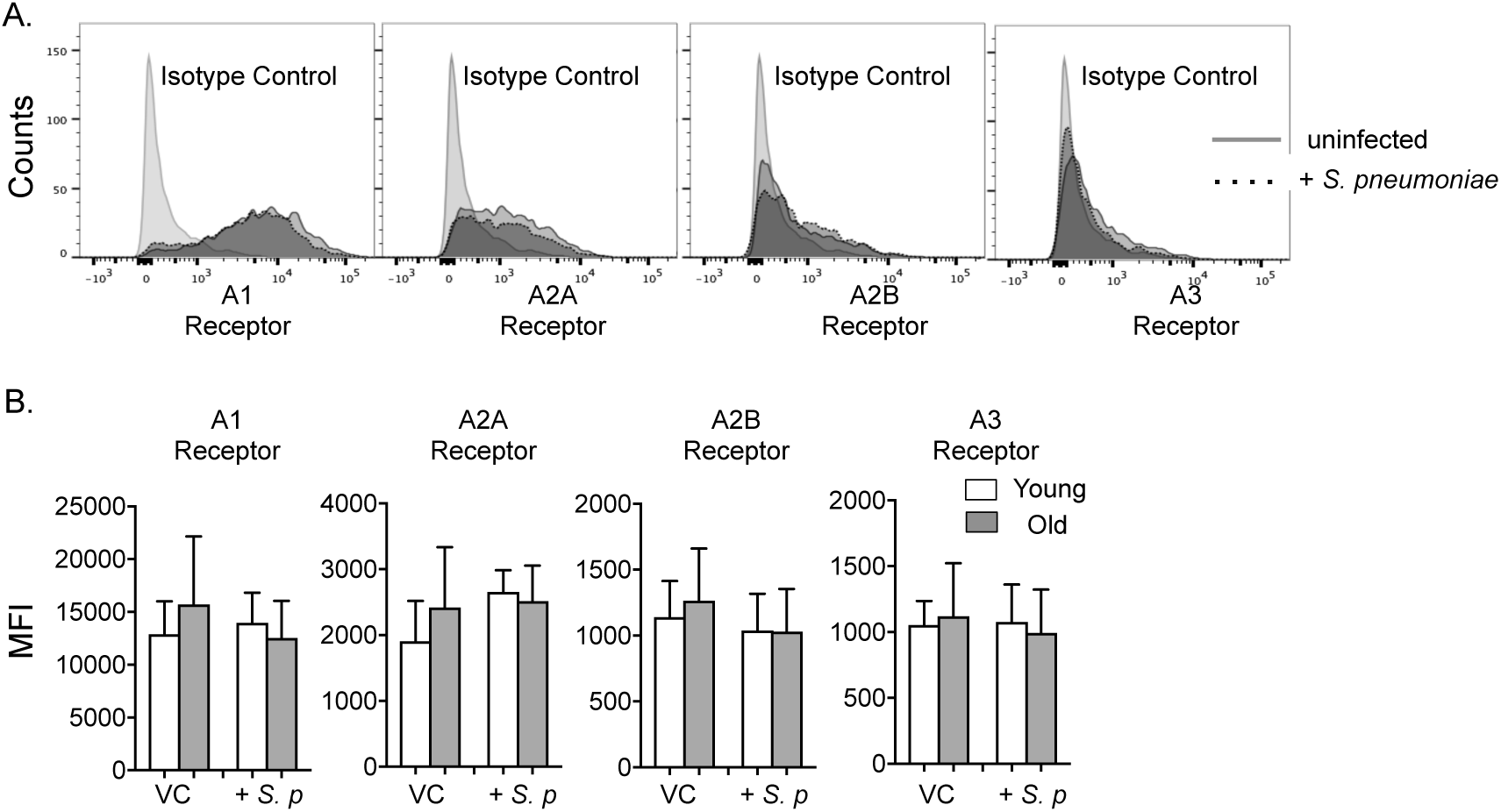
Pulmonary expression of adenosine receptors in mice are not altered by aging or *S. pneumoniae* infection. Young and old C57BL/6J mice were infected with *S. pneumoniae* intra-tracheally. The lungs were harvested six hours post infection and digested into a single cell suspension and the expression of adenosine receptors analyzed by flow cytometry. (A) Histograms shown are representative data from young mice and (B) bar graphs are quantification of the Mean Florescent Intensities (MFI) of the indicated adenosine receptor from young and old mice and are pooled from six separate experiments with n=9-12 mice per group.

We then evaluated the role of A1 receptor signaling in host resistance to infection. We treated mice with i.p. injections of DPCPX, a A1 pharmacological inhibitor that is active *in vivo* prior to infection. The mice were then challenged intra-tracheally with ∼1×10^4^ CFU, a low non-lethal dose of *S*. *pneumoniae* (Bou Ghanem, Clark, Du, et al., 2015; Bou Ghanem, Clark, Roggensack, et al., 2015), and lung bacterial burdens were determined 6 hours post-infection. Since extracellular adenosine modulates host resistance to infection via regulating the inflammatory immune response following infection (Aeffner et al., 2014, 2015; Barletta, Cagnina, et al., 2012; Barletta, Ley, et al., 2012; Bou Ghanem, Clark, Roggensack, et al., 2015; Theatre et al., 2012), we chose to challenge mice with this low dose of bacteria and examine bacterial numbers in the lungs at this early time point, conditions under which we previously found overt pulmonary inflammation is not yet triggered (Bou Ghanem, Clark, Roggensack, et al., 2015). We further confirmed that the A1 inhibitor had no effect on the baseline numbers of neutrophils, macrophages and T cells in the lungs (data not shown). We found that inhibition of A1 receptor signaling *in vivo* resulted in a significant ∼10-fold increase in bacterial numbers in the lungs of young mice following pulmonary challenge (Fig 8A). At 6 hours post infection, the infection was still localized to the lungs and had not spread systemically. However, when we compared bacteremia at 24 hours following infection, we found that while none of the control mice were bacteremic, 25% of A1 inhibited mice had systemic spread of the infection (Fig 8B). Importantly, inhibition of A1 resulted in significantly more host death when mice were challenged at an otherwise non-lethal dose (Fig 8D). These findings demonstrate that specific signaling via the A1 receptor is required for early host resistance to *S. pneumoniae* lung infection.

**Fig 8.**
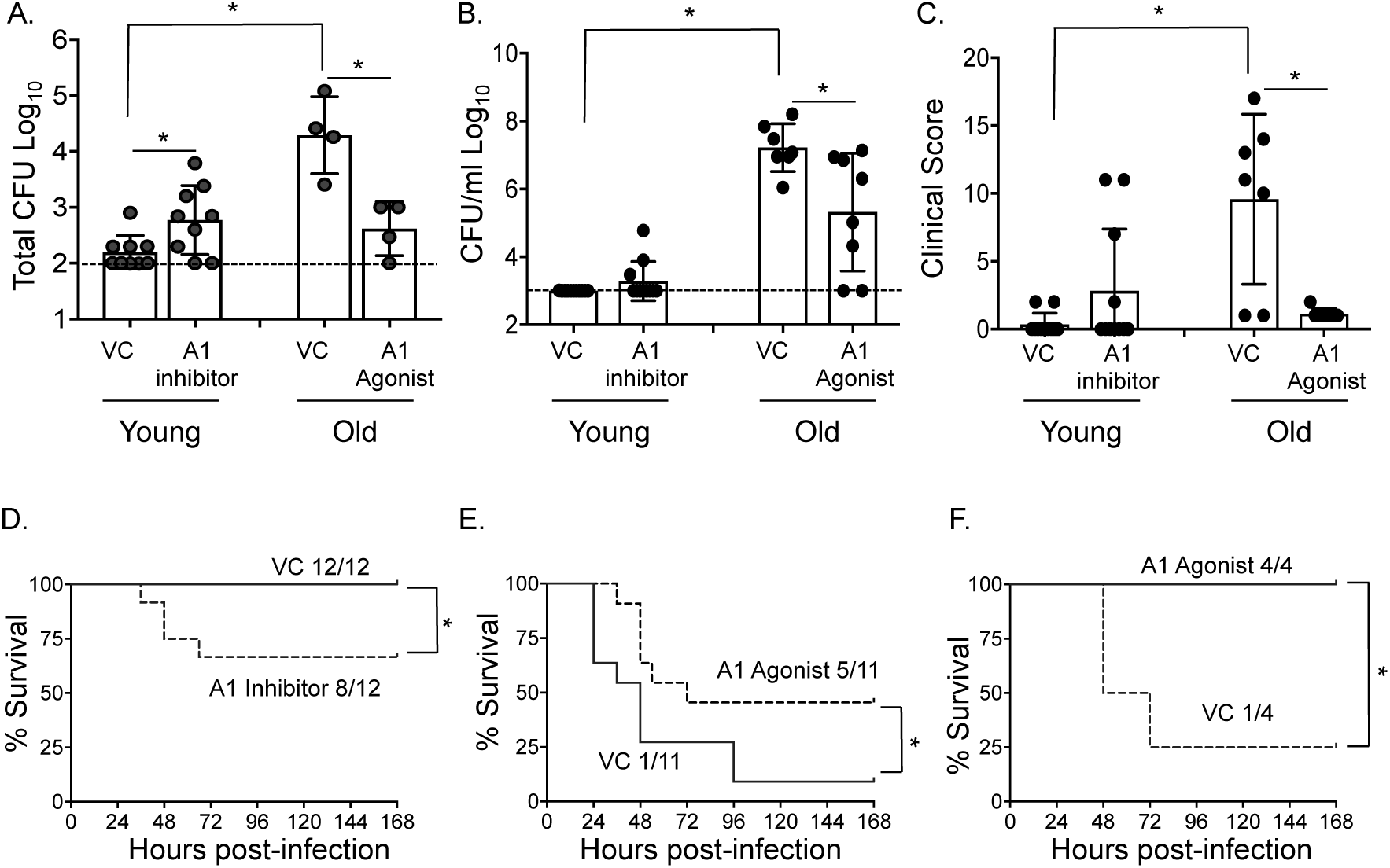
Signaling via A1 receptor is important for the control of pulmonary pneumococcal burden *in vivo.* Young mice mock-treated or treated with DPCPX, an A1 receptor inhibitor and old mice mock-treated or treated with 2-Chloro-N6-cyclopentyladenosine, an A1 receptor agonist were inoculated intra-tracheally with ∼1-2×10^4^ CFU of *S*. *pneumoniae* TIGR4. (A) Bacterial burdens in the lungs were determined 6 hours post-infection. Data shown are pooled from n=9 young and n=4 old mice per group. (B-E) To follow the infection over time, mice were challenged with ∼1-2×10^4^ CFU of *S*. *pneumoniae* TIGR4. Bacterial burdens in the blood (B) and clinical scores (C) were determined 24 hours post-infection. (A-B) Dashed lines indicate the limit of detection. Asterisks indicate significant differences calculated by one-way ANOVA followed by Tukey’s test. Survival of control and A1 inhibited young mice (D) as well as control and A1 agonist-treated old mice (E) and was assessed at the indicated time points following challenge. (F) Young mice control treated or administered A1 receptor agonist were infected with ∼1-2×10^5^ CFU, a lethal dose of *S*. *pneumoniae* TIGR4 and survival was followed over time. Fractions denote survivors at 7 days after challenge over the total number of mice. Asterisks indicate a statistical significance by the log-Rank (Mantel-Cox) test. Data were pooled from two independent experiments.

### Triggering A1 receptor signaling boosts resistance against S. pneumoniae lung infection in aged hosts

As pneumococcal infections are particularly a problem in the elderly, we examined the effect of aging on adenosine receptor expressions in the lung. We compared pulmonary expression levels of the adenosine receptors in young (2 months) versus old (18-22 months) mice. We found that there were no significant differences in the expression of any of the adenosine receptors in the lungs of old mice as compared to young controls either at baseline or following lung challenge with *S. pneumoniae* (Fig 7B).

We previously found that mice mimic the age-driven susceptibility to *S. pneumoniae* lung infection observed in humans (Bou Ghanem, Clark, Du, et al., 2015). To test whether A1 receptor can boost the resistance of old mice to lung infection, mice were treated i.p. with 2-Chloro-N6-cyclopentyladenosine, the A1 agonist prior to infection. Mice were then challenged intra-tracheally with *S*. *pneumoniae*, and bacterial burdens in the lung were determined 6 hours post-infection. As expected, vehicle control treated old mice were more susceptible to pneumococcal infection and had ∼ 100-fold more bacteria in their lungs at 6 hours (Fig 8A) and blood at 24 hours (Fig 8B) as compared to young controls.

Strikingly, treatment with the A1 agonist boosted the resistance of old mice to infection. A1 agonist treated old mice had 70-fold less bacteria in their lungs, a bacterial burden that was indistinguishable from that of control treated young mice (Fig 8A). Further, by 24 hours post infection, old mice given the A1 agonist had significantly less bacteria in their systemic circulation when compared to controls (Fig 8B). Importantly, triggering A1 significantly decreased clinical sign of sickness (Fig 8C) and boosted survival of old mice (Fig 8E). We also confirmed that treatment with the A1 agonist boosted survival of young mice that were challenged with a lethal bacterial dose (Fig 8F). These data demonstrate that A1 receptor can be targeted to boost the innate resistance of aged hosts to *S. pneumoniae* lung infection.

## Discussion

The role of the extracellular adenosine pathway in modulating host resistance against pulmonary pathogens is becoming better appreciated. Previous studies using murine models of infection showed that deficiency of adenosine A2B or A1 receptors was protective against *Klebsiella pneumoniae* (Barletta, Cagnina, et al., 2012) and influenza (Aeffner et al., 2014) lung infection respectively, whereas overexpression of CD39 on airway epithelial cells promoted *Pseudomonas aeruginosa* clearance from the lungs (Theatre et al., 2012). We previously found that extracellular adenosine production by CD73 was crucial for control of pulmonary bacterial numbers and systemic spread following lung challenge with *S. pneumoniae* (Bou Ghanem, Clark, Roggensack, et al., 2015). In all of the above studies, extracellular adenosine was believed to modulate pathogen clearance by either controlling the recruitment of immune cells, particularly neutrophils and/or regulating their antimicrobial activity (Aeffner et al., 2014, 2015; Barletta, Cagnina, et al., 2012; Barletta, Ley, et al., 2012; Bou Ghanem, Clark, Roggensack, et al., 2015; Theatre et al., 2012). Here we show that apart from its effect on innate immune cells, extracellular adenosine modulates host resistance by acting on pulmonary epithelial cells, targeting expression of mammalian proteins co-opted by *S. pneumoniae* as receptors and reducing the ability of bacteria to bind host cells.

Few studies have examined the effect of adenosine on bacterial infection of epithelial cells from different organs (Crane & Shulgina, 2009; Kao et al., 2017; Pettengill, Lam, & Ojcius, 2009; Spooner, DeGuzman, Lee, & Yilmaz, 2014; Walia et al., 2004). Adenosine signaling via the A2B receptor inhibited intracellular persistence of *Chlamydia trachomatis* in cervical epithelial cells (Pettengill et al., 2009) while A2A receptor promoted intracellular replication of *Porphyromonas gingivalis* in gingival epithelial cells (Spooner et al., 2014). In intestinal epithelial cells, adenosine promoted bacterial infections, where addition of exogenous adenosine increased adhesion of *Salmonella* to gut epithelial cells (Walia et al., 2004), and expression of the adenosine producing enzyme CD73 on intestinal epithelial cells promoted the intracellular replication as well as translocation and systemic spread of *Salmonella* across the gut epithelium (Kao et al., 2017). Similarly, adenosine altered the adhesion patterns of enteropathogenic *Escherichia coli* to gut epithelial cells and promoted intestinal infection *in vivo* (Crane & Shulgina, 2009). Thus the effect of adenosine on bacterial infection of the epithelium may be organ and adenosine-receptor specific. We demonstrated here that adenosine signaling via the A1 receptor blinted adherence of *S. pneumoniae* to H292 epithelial cells and reduced lung infection *in vivo*, which is to our knowledge, the first report to show an effect of the extracellular adenosine pathway on bacterial infection of pulmonary epithelial cells. In certain infections adenosine had a direct effect on bacterial viability and expression of virulence genes (Crane & Shulgina, 2009; Kao et al., 2017). However, that was not the case here as adenosine neither affected *S. pneumoniae* viability or expression of capsular polysaccharide, a major determinate of bacterial adhesion, nor did treatment of the bacteria alone alter their ability to adhere to lung epithelial cells.

Purinergic signaling plays a crucial role in regulating responses in the airways (Burnstock et al., 2012). We found here that adenosine was exerting its effect on bacterial binding through signaling via adenosine receptor A1 on host epithelial cells. The pulmonary epithelial cells used *in vitro* in this study were H292 human lung epithelial cells that are a mix of type I and type II pneumocytes (Bhowmick et al., 2013). We found that H292 cells expressed mRNA for A2B, A1 and A2A in decreasing levels while A3 levels were extremely low. This is consistent with previous studies with A549 cells, a type II pneumocyte cell line (Roman et al., 2006) as well as primary bronchial epithelial cells (Zhong et al., 2006). Similar to other studies, despite differences at the mRNA levels, we could detect protein expression for all four adenosine receptors (Giacomelli et al., 2018). Lung epithelial cells are also known to express P2 purinergic receptors that respond to ADP and/or ATP (Burnstock et al., 2012) and ATP and/or ADP impaired replication of *Chlamydia trachomatis* in cervical epithelial cells (Pettengill et al., 2012). However, addition of ATP did not alter *S. pneumoniae* binding to the lung epithelial cells *in vitro* while AMP significantly decreased bacterial binding. We believe that AMP but not ATP decreased bacterial binding since H292 cells express CD73 that can dephosphorylate AMP into adenosine but lack CD39 that is required for converting ATP into AMP. *In vivo*, the ability of epithelial cells to produce adenosine themselves may not be relevant as adenosine can be produced by many cells that influx into sites of infection and damage including immune cells (Eltzschig, Macmanus, & Colgan, 2008).

Here, A1 receptor signaling was important for control of *S. pneumoniae* lung infection *in vivo*. We found that A1 and A2A receptors were expressed in mouse lungs, while expression of A2B and A3 was lower, similar to what has been described before (Factor et al., 2007). A1 receptor is known to play an important role in acute lung injury by regulating inflammation, particularly pulmonary influx of neutrophils (Aeffner et al., 2014; Fernandez, Sharma, LaPar, Kron, & Laubach, 2013; Ngamsri, Wagner, Vollmer, Stark, & Reutershan, 2010; Sun et al., 2005). While we cannot differentiate whether the effect of adenosine *in vivo* is due to a direct effect on the lung epithelium or innate immune cells, we used a low dose infection and examined bacterial burdens early on after 6 hours of infection since we previously found that at this dose and this time point very little PMNs influx into the airways (Bou Ghanem, Clark, Roggensack, et al., 2015).

We found that adenosine and A1 receptor signaling down regulated expression of PAFR on *S. pneumoniae* infected lung epithelial cells. PAFR is a host protein used by *S. pneumoniae* to adhere to airway epithelial cells (Cundell, Gerard, et al., 1995). When PAFR was inhibited, we found that bacterial binding to the epithelium was reduced by half, consistent with previous reports (Cundell, Gerard, et al., 1995; Shukla et al., 2016). Triggering A1 receptor did not further reduce binding when PAFR was inhibited, suggesting that signaling via A1 receptor reduces pneumococcal binding via targeting PAFR. PAFR is expressed in the lungs (Duitman et al., 2012) and impairs host resistance against pneumococcal pneumonia (Cundell, Gerard, et al., 1995; Iovino, Brouwer, van de Beek, Molema, & Bijlsma, 2013; Rijneveld et al., 2004). We observed that PAFR expression is up regulated on lung epithelial cells upon infection, consistent with previous reports (Duitman et al., 2012). Therefore the reduced expression of PAFR in infected cells treated with adenosine or the A1 receptor agonist may be an indirect effect of less bacteria bound, however, we believe that this is unlikely to be a simple reflection of bound bacteria since expression of both ICAM-1 and PIGR, two other host proteins up regulated upon infection were not affected by adenosine. It is more likely that adenosine is indirectly regulating PAFR expression by controlling inflammatory responses of epithelial cells to infection. Previous studies found that PAFR was up regulated on epithelial cells activated by cytokines or other factors, which enhances bacterial binding (Cundell, Gerard, et al., 1995; Ishizuka et al., 2001; Mushtaq et al., 2011), and adenosine receptor signaling controlled cytokine responses by epithelial cells (Zhong et al., 2006). This could explain why we only see A1 receptor signaling reducing expression of PAFR in *S. pneumoniae* infected but not resting epithelial cells. In addition to bacterial adherence to the lung epithelium, PAFR also has a role in pneumococcal invasion of epithelial and endothelial cells and systemic spread of the infection from the lungs to the circulation *in vivo* (Cundell, Gerard, et al., 1995; Fillon et al., 2006; Radin et al., 2005). However, we did not detect any bacteremia in either A1 inhibited or control treated mice at the early 6-hour time point following *S. pneumoniae* lung challenge.

Changes in extracellular adenosine production, metabolism and signaling occur with aging (Chen & Chern, 2011; Headrick, 1996; Willems, Ashton, & Headrick, 2005). This is reflected by changed levels and activity of the EAD-producing and degrading enzymes (Crosti et al., 1987; Mackiewicz et al., 2006), altered expression of adenosine receptors (Willems et al., 2005), and an overall decline in G-protein coupled receptor signaling (Yeo & Park, 2002). This pathway has been reported to play an important role in the age-related decline of brain (Mackiewicz et al., 2006), metabolic (Rolband, Furth, Staddon, Rogus, & Goldberg, 1990) and cardiac function (Willems et al., 2005), and importantly, can be targeted to reverse this decline in animal models (Headrick et al., 2003; Hesdorffer et al., 2012; Reaven, Chang, & Hoffman, 1989). However, the role of the extracellular adenosine pathway in age-driven susceptibility to infections remains completely unexplored. We found here that there was no difference in the expression levels of adenosine receptors in the lungs of young versus old mice. Strikingly, treatment of old mice with an A1 receptor agonist significantly boosted their resistance to pneumococcal lung infection where they were able to control early pulmonary bacterial burdens, had reduced systemic spread of the infection and extended survival compared to control treated mice. PAFR expression was reported to be upregulated at baseline in the lungs of old mice (Shivshankar et al.). Therefore it is possible that triggering A1 receptors boots the resistance of aged hosts to *S. pneumoniae* by reducing PAFR expression in the lungs. These findings demonstrate that targeting the adenosine pathway may be a feasible strategy to reverse the innate susceptibility of aging to lung infection.

*S. pneumoniae* binds to PAFR via phosphorylcholine displayed on the bacterial cell wall that mimics moieties found on host platelet-activating factor. This form of molecular mimicry is shared by other pathogens (S. E. Clark & Weiser, 2013), including *Haemophilus influenzae* (Swords et al., 2000), *Neisseria* (Serino & Virji, 2002; Weiser, Goldberg, Pan, Wilson, & Virji, 1998), *Pseudomonas aeruginosa* (Barbier et al., 2008)*, Acinetobacter baumannii* (Smani et al., 2012) and is used by bacteria to bind to PAFR and adhere to host cells but can also regulate inflammatory responses in the host (S. E. Clark & Weiser, 2013; Hergott et al., 2015). Intriguingly a pilot study found that single nucleotide polymorphisms in *PTAFR* gene coding for PAFR was associated with increased risk of invasive pneumococcal disease in humans, although these results need to be validated as no correction for multiple testing was performed (Lingappa et al., 2011). In conclusion, our findings regarding the effect of adenosine on PAFR expression and bacterial binding on lung epithelial cells may have future implications for using clinically available adenosine-based therapies (Hasko et al., 2008) to boost host resistance to *S. pneumoniae* and other serious lung infections.

## Experimental Procedures

### Mice

Young (2 months) and Old (18-22 months) C57BL/6 mice were purchased from Jackson Laboratories (Bar Harbor, ME) and the National Institute on Aging colonies and housed in a specific-pathogen free facility at the University at Buffalo. Male mice were used in all experiments. This work was performed in accordance with the recommendations in the Guide for the Care and Use of Laboratory Animals published by the National Institutes of Health. All procedures were reviewed and approved by the University at Buffalo Institutional Animal Care and Use Committee.

### Bacteria

Wild type *S. pneumoniae* TIGR4 strain (serotype 4), were grown at 37°C with 5% CO_2_ in Todd-Hewitt broth (BD Biosciences) supplemented with 0.5% yeast extract and Oxyrase (Oxyrase) untill mid-exponential phase. Aliquots were then frozen at -80°C in the growth media with 25% (v/v) glycerol. Before use, bacterial aliquots were thawed on ice, washed once and diluted in PBS to the required concentrations. Bacterial titers were confirmed by plating on Tryptic Soy Agar plates supplemented with 5% sheep blood agar (blood agar plates-Northeast Laboratory Services). The Δ*cps* deletion mutant was a kind gift from Andrew Camilli (Shainheit, Mule, & Camilli, 2014).

### PMN isolation

PMNs were isolated from the peripheral blood of healthy donors as previously described (Bou Ghanem et al., 2017). Volunteers were recruited in accordance with the University at Buffalo Human Investigation Review Board (IRB) and signed informed consent forms.

### Bacterial binding assay with H292 cells

Human pulmonary mucopidermoid carcinoma-derived NCI-H292 (H292) cells were purchased from ATCC and grown at 37°C/ CO_2_ in “H292” media consisting of RPMI 1640 medium (ATCC) with 2 mM L-glutamine, 10% FBS, and 100 U penicillin/streptomycin following a previously described (Bhowmick et al., 2013). The day before the binding assay 2.5 x 10^5^ H292 cells were seeded in tissue culture-treated flat bottom 96-well plates (Corning) and allowed to adhere overnight. The following day, cells were washed in PBS and antibiotic-free H292 media was added. The cells were then pretreated for 30 minutes with the indicated drugs or PBS as a vehicle control and then infected with *S. pneumoniae* at an MOI of 10-20. The plates were spun down and incubated for 1 hour at 37°C/ CO_2_. To determine the amount of bacteria that adhered, the cells were washed with PBS, lifted with 0.05% Trypsin/EDTA (Invitrogen) and vigorously vortexed to produce a homogeneous solution. Serial dilutions were then plated on blood agar plates for determination of bacterial colony forming units (CFU). The percent of bacteria bound were determined with respect to a no cell control where bacteria were added to the wells and incubated for an hour under the same experimental conditions (+/-drugs) and then viable bacteria enumerated by plating on blood agar plates. The amount of bacteria bound to control treated cells was set as 100 % and relative changes in bacterial binding in treated cells was then calculated and is presented as % bound relative to vehicle treated controls on the graphs. The effect of adenosine treatment on bacterial binding was also confirmed by performing binding assays on H292 cells seeded on glass slides and lysed with water (not shown).

### Adenosine pathway and PAFR drugs

The effect of the extracellular adenosine pathway was tested using the following: Adenosine (Sigma) diluted in PBS and used at the concentrations indicated on the graphs. The pan adenosine receptor agonist 5′-(N-Ethylcarboxamido) adenosine (NECA) used at 1µM. This agonist targets all four receptor with reported Ki values of 6.2, 14, and 20 nM for human A3, A1 and A2A receptors respectively and EC50 of 2.4 µM for human A2B. The pan adenosine receptor inhibitor (ADRI) CGS-15943 used at 1µM. This inhibitor targets all four adenosine receptors, with Ki values of 3.5, 4.2, 16 and 51 nM for human A1, A2A, A2B and A3 receptors respectively. The A1 agonist 2-Chloro-N6-cyclopentyladenosine used at 10, 1, 0.1 nM, A2A agonist CGS21680 used at 270, 27, 2.7nM and A2B agonist BAY 60-6583 used at 200, 20, 2nM. These concentrations correspond to the approximate 10xKi, Ki or 0.1Ki for each receptor. Adenosine receptor drugs were purchased from Tocris or Sigma Aldrich, dissolved in DMSO and dilutions performed in PBS. The potent PAFR antagonist Apafant was also purchased from Tocris and used at 40nM corresponding to 2xKi.

### Mouse infections

The effect A1 receptor on lung infection was determined using the competitive inhibitor of A1 receptor, 8-Cyclopentyl-1,3-dipropylxanthine (DPCPX) or A1 agonist 2-Chloro-N6-cyclopentyladenosine. All chemicals were purchased from Sigma Aldrich, dissolved in DMSO and filter sterilized by passing through a 0.22µm filter. The mice were then given intraperitoneal (i.p.) injections of 1mg/kg of the A1 inhibitor or 0.1mg/Kg of the A1 agonist or vehicle control at days -1 and 0 (immediately before challenge) relative to infection. Control mice were mock-treated with the vehicle control. Mice were challenged intra-tracheally with 1-2×10^4^ *S*. *pneumoniae* TIGR4 as previously described (Bou Ghanem, Clark, Roggensack, et al., 2015). Six hours post-infection, mice were euthanized and lung and blood samples were harvested and plated on blood agar plates for enumeration of bacterial loads. Following the infection, mice were monitored daily and scored blindly for signs of sickness including weight loss, activity, posture and breathing. Based on these criteria, the mice were given a clinical score of healthy [0] to severely sick [21] as previously described (Mook-Kanamori, Geldhoff, Troost, van der Poll, & van de Beek, 2012).

### Flow cytometry

For examining adenosine receptor expression in the lungs, mice were perfused with 10ml PBS, the lungs harvested and minced into small pieces. The lungs were then digested for 1 hour with RPMI 1640 supplemented with 10% FBS, 1 mg/ml Type II collagenase (Worthington), and 50 U/ml Deoxyribonuclease I (Worthington) at 37°C/ 5% CO_2_. Single-cell suspensions were obtained by mashing the digested lungs, and the red blood cells were removed by treatment with a hypotonic lysis buffer (Lonza). For examining expression of host proteins on H292 cells, the cells were infected with *S*. *pneumoniae* at an MOI of 2 as described in the binding assay. The cells were resuspended in FACS buffer (HBSS/ 1% FBS) then treated with Fc block (anti-mouse clone 2.4G2 and anti-human clone 3G8) purchased from BD and stained with specific antibodies. The following conjugated anti-human antibodies were used: PAFR (040582; Cederlane), ICAM (clone HA58; Biolegend), CD73 (clone TY/11.8; eBioscience), CD39 (clone24DMS-1; eBioscience) and PIGR (PA535340; Invitrogen). For staining for adenosine receptor antibodies, the cells were permeabilized using the BD Cytofix/Cytoperm kit. The following unconjugated primary rabbit polyclonal anti-adenosine receptor antibodies were purchased from abcam and used: anti-mouse and human A2a (ab3461), A2b (ab222901) and A3 (ab203298) as well as anti-mouse A1 (ab82477). Rabbit polyclonal IgG (ab37415) purchased from abcam and used as an isotype control. For anti-human A1 a primary rabbit monoclonal antibody was used (clone ERP6 179, ab124780). Secondary PE-conjugated anti-Rabbit IgG was used (12473981; Invitrogen). Fluorescence intensities were measured on a BD FACS Fortessa and data were analyzed using FlowJo.

### RNA extraction and qPCR

RNA was extracted from 2×10^5^ H292 using the RNeasy Mini Kit (Qiagen) as per manufacturer’s protocol. TURBO DNA-free kit (Invitrogen) was used to digest DNA from the RNA samples prior conversion into cDNA. For each sample, 500ng of RNA was converted into cDNA using SuperScript VILO^TM^ cDNA synthesis kit (Life Technologies) according to the manufacturer’s protocol. RT-PCR was performed using CFX96 Touch™ Real-Time PCR Detection System from Bio-Rad and CT (cycle thresh-hold) values were determined using the following TaqMan probes from Life Technologies (Thermo Fischer Scientific): GAPDH (Hs99999905_m1), A1 (Hs00181231_m1), A2A (Hs00169123_m1), A3 (Hs00252933_m1), and A2B (Hs00386497_m1). All samples were run in duplicates. Data were analyzed by the comparative threshold cycle (2^-ΔCT^) method, normalizing the CT values obtained for target gene expression to those for GAPDH of the same sample. For comparison of adenosine receptor levels upon infection, relative quantity of transcripts (RQ) values were calculated by the ΔΔCT method by using the formula RQ =2^^_^ (ΔΔCT) (Livak & Schmittgen, 2001). The ΔΔCT values were obtained by subtracting ΔCT value of the infected from that of the uninfected control.

### Western blotting

H292 cells were solubilized in radioimmunoprecipitation assay (RIPA) buffer (1% Triton X-100, 0.25% sodium deoxycholate, 0.05% SDS, 50 mM Tris-HCl [pH 7.5], 2 mM EDTA, 150 mM NaCl, 1 mM phenylmethylsulfonyl fluoride, 1 mM sodium orthovanadate, 10 mg/liter each of aprotinin and leupeptin). Lysates were centrifuged and protein concentrations of the supernatants were quantified using bicinchoninic acid kit (Pierce). Equal protein quantities of each sample were migrated on Mini-PROTEAN TGX Stain-Free Precast Gels (BioRad) and transferred to polyvinylidene difluoride (PVDF) membranes. Incubations with primary antibodies at 1:1000 dilutions were done overnight at 4°C. Incubation with secondary antibodies (1: 5000 dilutions) coupled to horseradish peroxidase was done for 1 hour at room temperature. ChemiDoc XRS+ system (BioRad) was used for detection of all Western blots using Clarity Western ECL Substrate (BioRad). Blots were stripped in stripping buffer containing glycine, Triton-X 100 and SDS, and probed for GAPDH, which served as a loading control. Primary antibodies against A1 (ab124780), A2A (ab3461), A2B (ab222901) and A3 (ab203298) were all obtained from Abcam. Primary antibodies against GAPDH (MA5-15738) and horseradish peroxidase-conjugated secondary antibodies (31460 and 31430) were purchased from Invitrogen. Images were quantified using ImageJ (version 1.51e) software. Integrated pixel densities in bands corresponding to proteins of interest were measured, and the background was subtracted. Integrated pixel densities for GAPDH as loading controls were also determined and corrected for the background. Background-corrected integrated pixel densities of the protein of interest were normalized to those of the loading control.

### Statistics

All statistical analysis was performed using Prism7 (Graph Pad). CFU data were log-transformed to normalize distribution. For all graphs, the mean values +/-SD are presented. Significant differences were determined by 1-sample t-test, Student’s t-test or one-way ANOVA followed by the Tukey’s multiple comparisons test as indicated. Survival analysis was performed using the log-rank (Mantel-Cox) test. *p* values less than 0.05 were considered significant (as indicated by asterisks).

## Acknowledgments

We would like to acknowledge Andrew Camilli for bacterial strains as well as Anthony Campagnari and Ira Blader for their feedback on the manuscript. Research reported in this publication was supported by the National Institute On Aging of the National Institutes of Health under Award Number R00AG051784. The content is solely the responsibility of the authors and does not necessarily represent the official views of the National Institutes of Health.

